# Variant scoring performance across selection regimes depends on variant-to-gene and gene-to-disease components

**DOI:** 10.1101/2024.09.17.613327

**Authors:** Hilary K. Finucane, Sophie Parsa, Jeremy Guez, Masahiro Kanai, F. Kyle Satterstrom, Lethukuthula L. Nkambule, Mark J. Daly, Cotton Seed, Konrad J. Karczewski

**Affiliations:** Program in Medical and Population Genetics, Broad Institute of MIT and Harvard, Cambridge, Massachusetts 02142, USA; Analytic and Translational Genetics Unit, Massachusetts General Hospital, Boston, Massachusetts 02114, USA; Center for Computational and Integrative Biology, Massachusetts General Hospital, Boston, Massachusetts, 02114, USA; Novo Nordisk Foundation Center for Genomic Mechanisms of Disease, Broad Institute of MIT and Harvard, Cambridge, Massachusetts 02142, USA

## Abstract

Variant scoring methods (VSMs) aid in the interpretation of coding mutations and their potential impact on health, but their evaluation in the context of human genetics applications remains inconsistent. Here, we describe GeneticsGym, a systematic approach to evaluating the real-world impact of VSMs on human genetic analysis. We show that the relative performance of VSMs varies across regimes of natural selection, and that both variant-to-gene and gene-to-disease components contribute.

## Main

Variant scoring methods (VSMs) such as PolyPhen2^1^ have been powerful tools for molecular diagnosis of rare disease and interpretation of association data for common disease. In recent years, many new variant scoring methods have been developed, each claiming greatly improved power. However, while each VSM presents benchmarks compared to previous VSMs, these benchmarks are often inconsistent, using different data and comparison methods. Moreover, the use of metrics such as AUC has made interpretation difficult, given the proposed applications are most often gene discovery or diagnostic practice. Independent benchmarking studies have used deep mutational scanning^2,3^, and shown concordance with clinical classification in ClinVar^4^, but these approaches primarily focus on protein-level effects and do not fully capture the complexities of variant impacts across a range of human diseases.

Here, we present a suite of evaluation metrics spanning a range of selective effects, including diseases ranging from severe developmental to common, late-onset disease, and variant allele frequencies ranging from *de novo* to common. We also use a large, independent population genetics data set to distinguish the ability of these metrics to identify the overall essentiality of a gene from the individual molecular impact of variants on a gene. We apply these evaluations to various VSMs^5–10^ and find that the best VSM depends on the specific disease and its selective regime.

We overcome two main challenges in constructing our evaluations. First, because most human genetic analyses focus on variants with the highest predicted pathogenicity, we avoid the use of AUC or Mann-Whitney U, both of which are largely driven by the relative rankings of variants with scores below the 90th percentile (Extended Data Fig. 1). Instead, we use statistics that quantify the amount of signal concentrated in the top 1-10% of variants: the rate ratio for analyses of *de novo* mutations, to maximize comparisons to previous work, and enrichment for analyses of other datasets. Second, as in previous benchmarking work^3^, we aggregate pairwise comparisons to avoid restricting to the small set of variants scored by all VSMs (Supplementary Information).

In our first evaluation, we assess the ability to distinguish *de novo* mutations in individuals with severe disorders compared to unaffected individuals in three trio-based cohorts (undiagnosed severe developmental disorders^11^, autism^12^, and congenital heart disease^13^; see Supplementary Information). The analysis gives a direct assessment of disease relevance without the risk of double-dipping, focusing on variants with extremely high selection coefficients. Analogous evaluations have been performed in previous papers, but none bring together the most recently available VSMs, and some use AUC or Mann-Whitney U. The two best-performing VSMs in our analysis are popEVE and MisFit *s* (Fig. 1a, Extended Data Fig. 2), although not all pairwise comparisons with the immediate runners up (MPC, AlphaMissense) are significant (Extended Data Fig. 3). Results for the three cohorts were similar (Extended Data Fig. 2), possibly due to sharing of controls and/or disease etiology. Both popEVE and MisFit *s* are based on a framework in which a within-gene score is calibrated across genes using population genetic data, suggesting that this is a successful framework for prioritizing variants in this regime of high selection.

**Figure 1.**
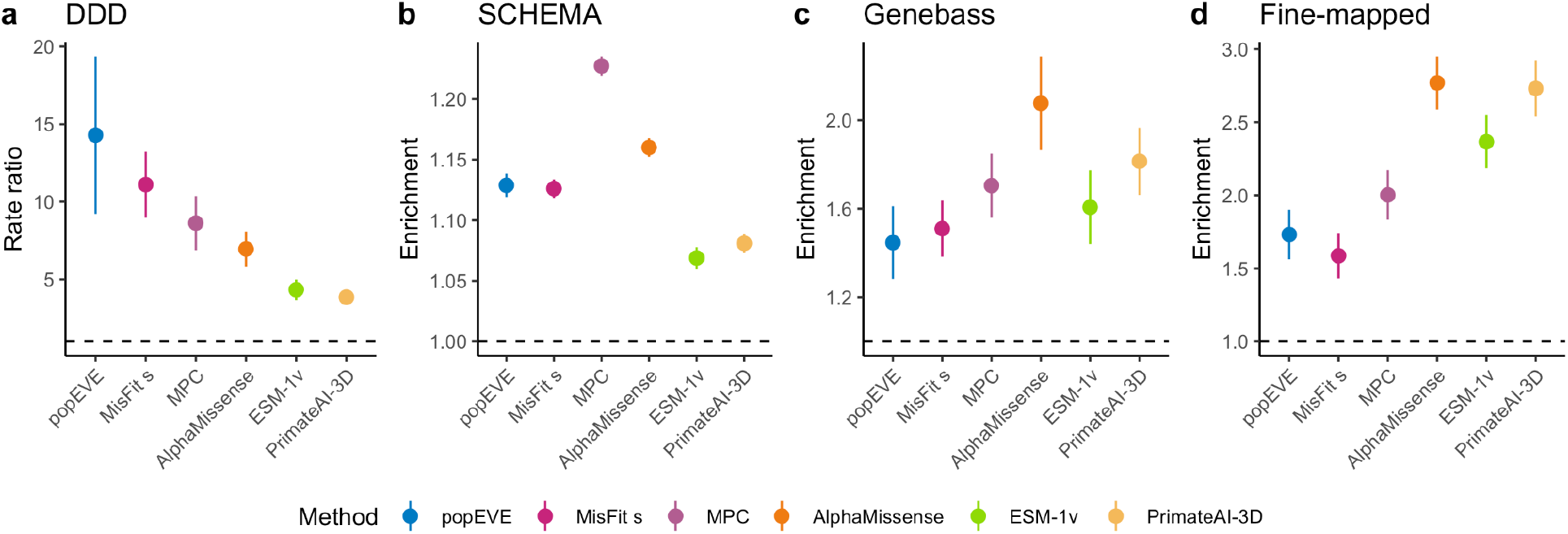
Rate ratios and enrichments for selected VSMs and selected evaluation datasets. (a) Rate ratio of *de novo* mutations in probands with severe undiagnosed developmental delay^11^ vs. unaffected siblings and controls. (b) Enrichment of rare variants (allele count < 30) in cases vs. controls from the Schizophrenia exome meta-analysis consortium (SCHEMA)^14^. (c) Enrichment of rare and low-frequency variants (allele frequency < 0.01) associated with a phenotype in the UK Biobank vs. frequency matched controls, from Genebass^15^. (d) Enrichment of fine-mapped variants from UK Biobank, FinnGen, and Biobank Japan vs frequency-matched controls^16^. In all panels, we show the rate ratio or enrichment based on a cutoff of the top 5% of all possible missense variants. Enrichments and rate ratios computed on different sets of variants are transformed to be comparable to each other using pairwise relative enrichments (see Supplementary Information). We show all evaluations, VSMs, and cutoffs in Extended Data Fig. 2, and significance of all pairwise comparisons in Extended Data Fig. 3.

Next, we assessed the ability of each VSM to identify rare variants conferring risk for schizophrenia, autism, and epilepsy. We used existing data from case-control exome sequencing studies of each of these disorders (N=24,248, 5,556, and 20,979, respectively) to compute an enrichment of variants found only in cases compared to variants found only in controls (Supplementary Information). Here, MPC was the highest performing VSM, followed by AlphaMissense (Fig. 1b, Extended Data Fig. 2). popEVE and MisFit *s* were no longer the top VSMs, likely because the variants identified in this study are mostly inherited variants and are not under as heavy selective pressure as the *de novo* variants contributing to severe disease.

The three disorders gave similar results (Extended Data Fig. 2), possibly due to partial sharing of controls and/or genetic similarities among the three diseases.

Moving to a regime in which selection is less strong, we assessed the ability of each VSM to identify rare and common variants contributing to common diseases and complex traits. To do this, we used rare variants with significant associations to at least one phenotype in the UK Biobank^15^, and common variant fine-mapping results (PIP > 0.1) in the UK Biobank, FinnGen, and Biobank Japan^16^ (Supplementary Information), against frequency-matched control variants. Because this evaluation focuses on common diseases and variants with higher allele frequencies than the other analyses, the relative performance of popEVE and MisFit *s* continued to drop, with AlphaMissense and PrimateAI-3D having highest performance (Fig. 1c,d, Extended Data Fig. 2).

The effect of a missense variant on disease risk reflects a combination of the effect of the variant on gene function and the effect of gene disruption on disease risk. To distinguish the extent to which VSMs are capturing these two components, we converted each VSM into a gene scoring method by assigning each variant the median score in the gene. We then evaluated each gene scoring method, in comparison to the overall performance of the VSM (Fig. 2a-d). We found wide variation in the extent to which each VSM excelled at gene scoring as opposed to variant scoring overall: for instance, MPC has the highest performance in identifying genes with high rate ratio/enrichment across all four evaluation datasets, while other methods provide variant-level insights including popEVE for DD and AlphaMissense for fine-mapped variants.

**Figure 2.**
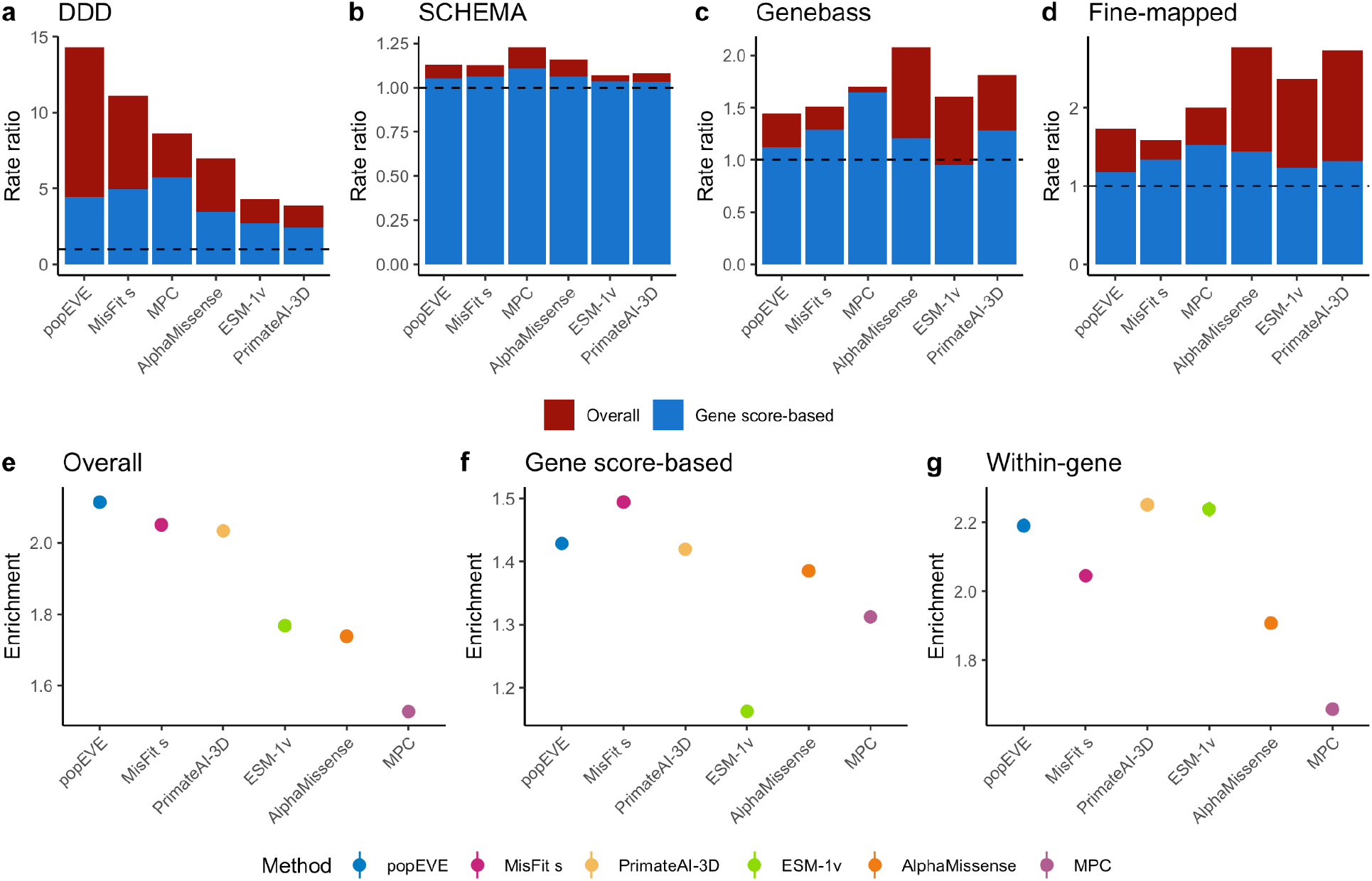
Influence of variant-to-gene and gene-to-disease scores on VSM performance. (a-d) Results for selected VSMs and evaluation sets as in Figure 1, with results for a gene score derived from the VSM by taking a median within each gene (gene score-based), and the raw results for the VSM (overall). (e) Enrichment of each VSM for identifying variants observed vs unobserved in gnomAD v4, restricting to sites at which variation was not observed in the training data of the VSMs benchmarked here (“gnomAD-new”). (f) Enrichment of each VSM gene score on gnomAD-new. As in (a-d), each VSM is transformed into a gene score by assigning to every variant the median score in its gene. (g) The average across genes of the enrichment of each VSM on gnomAD-new, computed separately for each gene. In all panels, we show the enrichment at the top 5% of variants. For (e-g), analyses are restricted to the set of 50,468,596 variants scored by all VSMs, and percentiles are computed on the set of variants scored by all VSMs with labels in gnomAD-new. Significance of all pairwise comparisons is shown in Extended Data Fig. 4. For panels e-g, scores were re-ordered by their performance in gnomAD-new (e).

To more precisely compare the extent to which different VSMs excel at ranking genes vs ranking variants within a gene, we used a large population genetics dataset, gnomAD v4, that provided sufficient power to assess both within-gene and across-gene predictions. To prevent double-dipping, we restricted our analyses to sites at which variation had not been observed in gnomAD v2 or the UK Biobank. Considering only those sites, we assessed the ability of each VSM to distinguish between variants observed in gnomAD v4 vs any possible missense variant that was not observed in gnomAD v4. This set of evaluation labels is independent of the data used for training any of the VSMs benchmarked here, and we call it gnomAD-new.

First, we used gnomAD-new to assess each VSM overall (Fig. 2e)l. As with the *de novo* variant evaluation, we found that popEVE and MisFit s were again the top performers, likely because this evaluation also focuses on the heavy selection regime. To assess the contribution of overall gene ranking to the overall performance of each VSM on gnomAD-new, we used gnomAD-new to assess each VSM-based to gene score, as above (Fig. 2f). Finally, we assessed the ability of each VSM to distinguish between observed and unobserved variants *within* a gene by performing separate analyses of each VSM within each gene and averaging the results across all genes (Fig. 2g), again using gnomAD-new labels. The top-performing VSMs in the gene score-based analysis (MisFit *s*, popEVE) were different from the top-performing VSMs in the within-gene analysis (PrimateAI-3D, ESM-1v). Overall performance is similarly correlated with gene score-based performance and within-gene performance, indicating that performance on both sub-tasks contributes to overall performance. These results suggest there is room for further improvement in variant effect prediction for this task, for example by combining the best-performing within-gene VSMs with the best-performing gene scores.

Finally, we assessed the extent to which the within-gene performance of these VSMs varies by gene type and function by repeating our within-gene performance evaluation (Fig. 2g) on subsets of genes (Extended Data Fig. 5). While we do not find ClinVar to be useful for overall benchmarking due to the risk of double-dipping, we include it in our comparison of accuracy across different gene sets (Extended Data Fig. 6). For both gnomAD-new and ClinVar, we find that, indeed, the relative performance of the VSMs changes depending on the gene set used for evaluation. For example, while all VSMs have higher performance in constrained genes compared to unconstrained genes, MisFit *s* shows substantially lower relative enrichments in constrained genes. While absolute enrichments using ClinVar are confounded by its use in hyperparameter tuning for AlphaMissense and likely implicit in many methods due to ClinVar benign variants found in population datasets used in training, a comparison of performance on recessive vs dominant genes shows that PrimateAI-3D has higher enrichment for recessive genes than dominant genes.

In summary, GeneticsGym provides a comprehensive evaluation of VSMs across multiple data sources and metrics, revealing that the best-performing VSMs vary depending on the specific use case and evaluation criteria. This highlights the importance of selecting appropriate predictors for different genetic contexts and research questions, and also the importance of benchmarking using varied data. Additionally, our work highlights the influence of variant-to-gene as well as gene-to-disease effects, which can lay groundwork for future scores that combine the best performing elements of each metric. We hope this resource will facilitate the development and refinement of these crucial tools, ultimately improving our ability to interpret genetic variants and their impact on human health.

## Acknowledgements

This work was supported by the Novo Nordisk Foundation (NNF21SA0072102).

## Extended Data

**Extended Data Figure 1.**
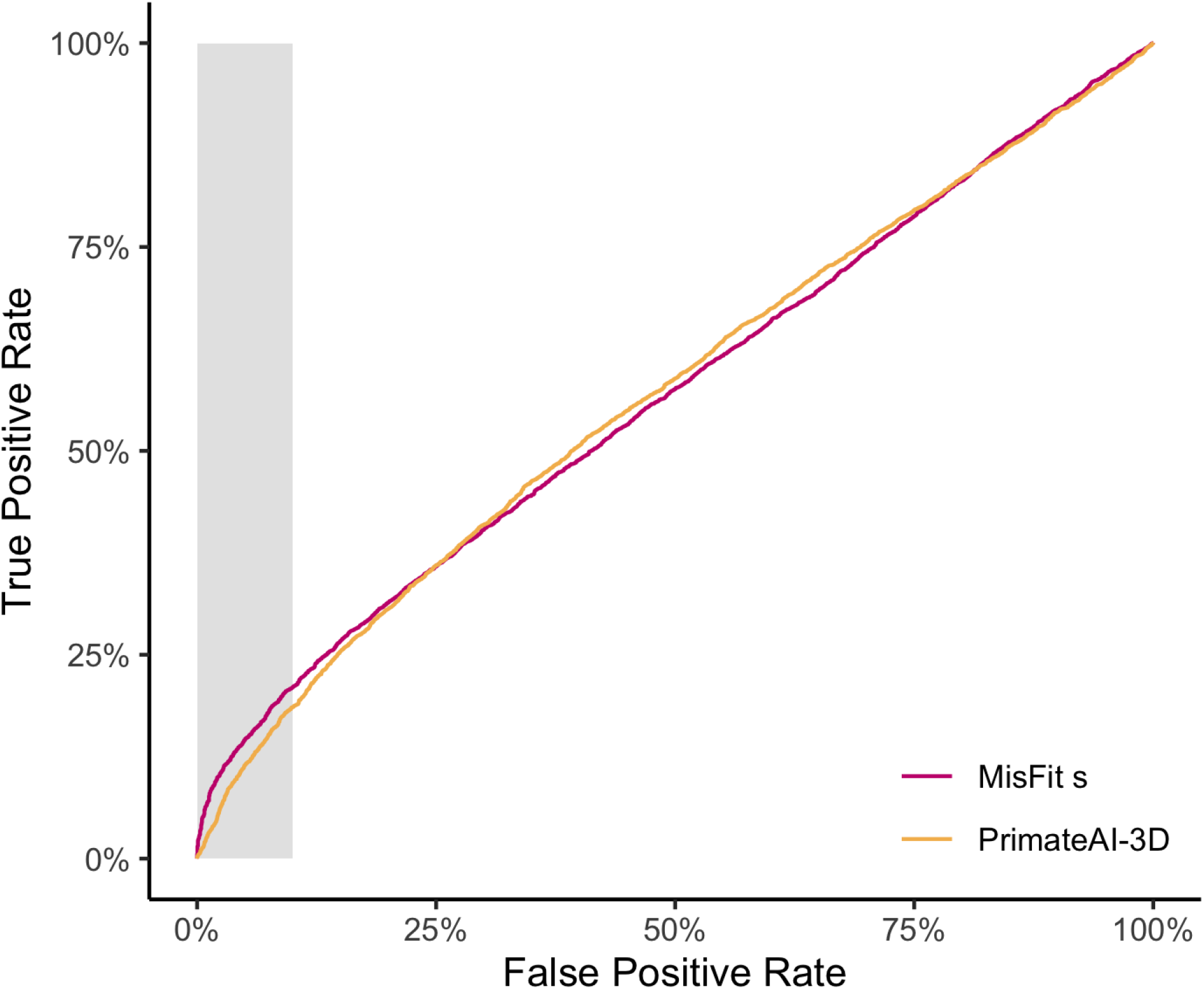
AUC can be a misleading method for comparisons with focused signal. We assessed the ability of VSMs to distinguish *de novo* variants in individuals with developmental disorders vs those in controls. The most informative regime of this plot is highlighted in gray; cutoffs with higher false positive rates will not often be useful for human genetics analyses. The higher true positive rate of MisFit *s* in the gray region indicates its higher utility in the most relevant regime. However, AUC is slightly higher for PrimateAI-3D (0.569) than MisFit s (0.568). AUC can thus be a misleading metric in cases such as this one, as it reflects the area under the entire curve, failing to focus on the most relevant area.

**Extended Data Figure 2.**
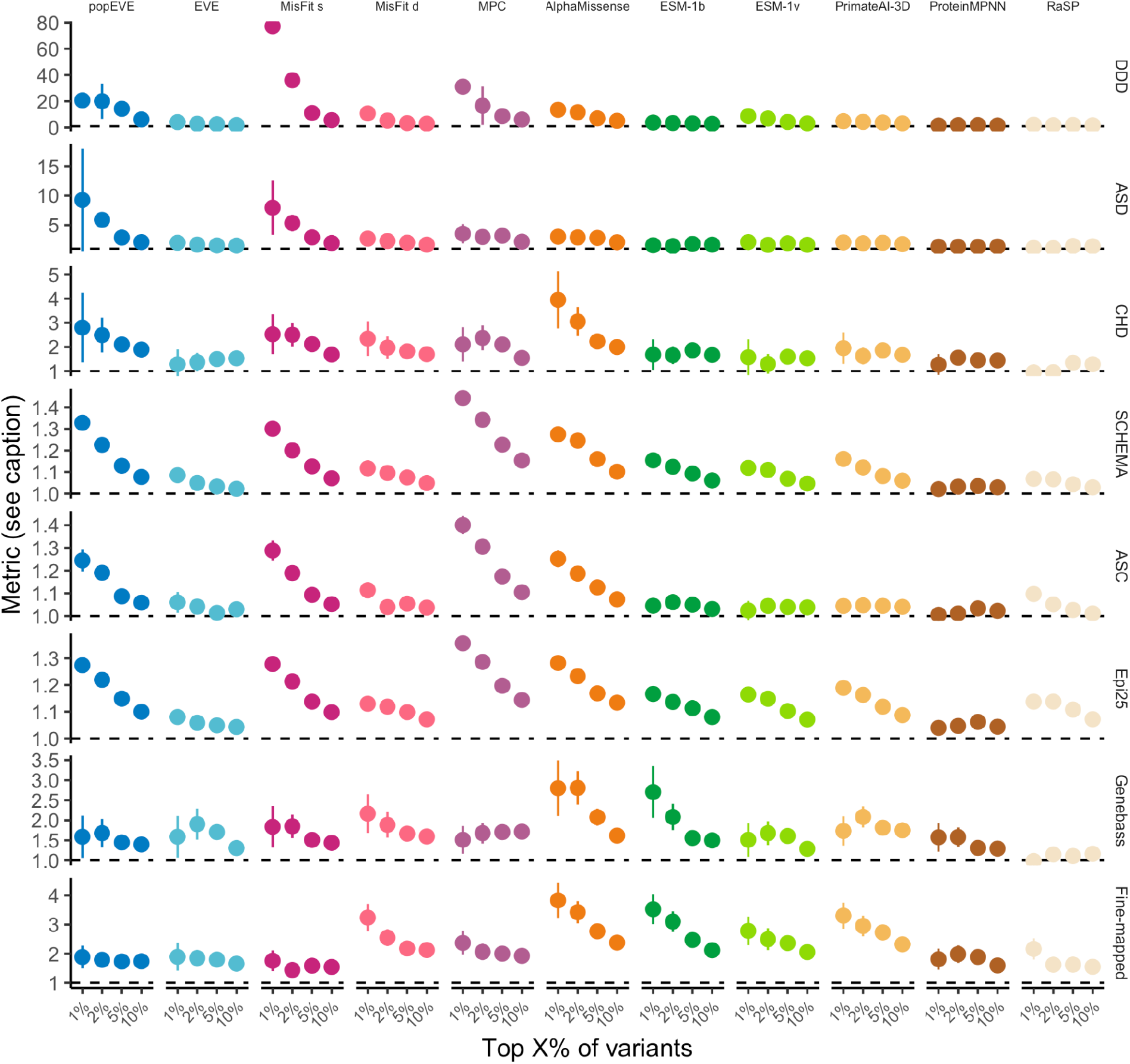
Evaluations of all scores and all evaluation data sets, at four cutoffs. Rows 1-3: Rate ratio of *de novo* mutations in probands vs. unaffected siblings, for all VSMs and four cutoffs. Data from developmental delay/intellectual disability (DD)^11^, autism (ASD)^12^, and congenital heart disease (CHD)^13^. Rows 4-6: Enrichment of rare variants in cases vs. controls from schizophrenia (SCHEMA)^14^, autism (ASC)^17^, and epilepsy (Epi25)^18^. Row 7: Enrichment of rare and low-frequency variants (allele frequency < 1%) associated with a phenotype in the UK Biobank vs. frequency matched controls, from Genebass^15^. Row 8: Enrichment of fine-mapped variants from UK Biobank, FinnGen, and Biobank Japan vs frequency-matched controls^16^. Enrichments and rate ratios computed on different sets of variants are transformed to be comparable to each other using pairwise relative enrichments (see Supplementary Information).

**Extended Data Figure 3.**
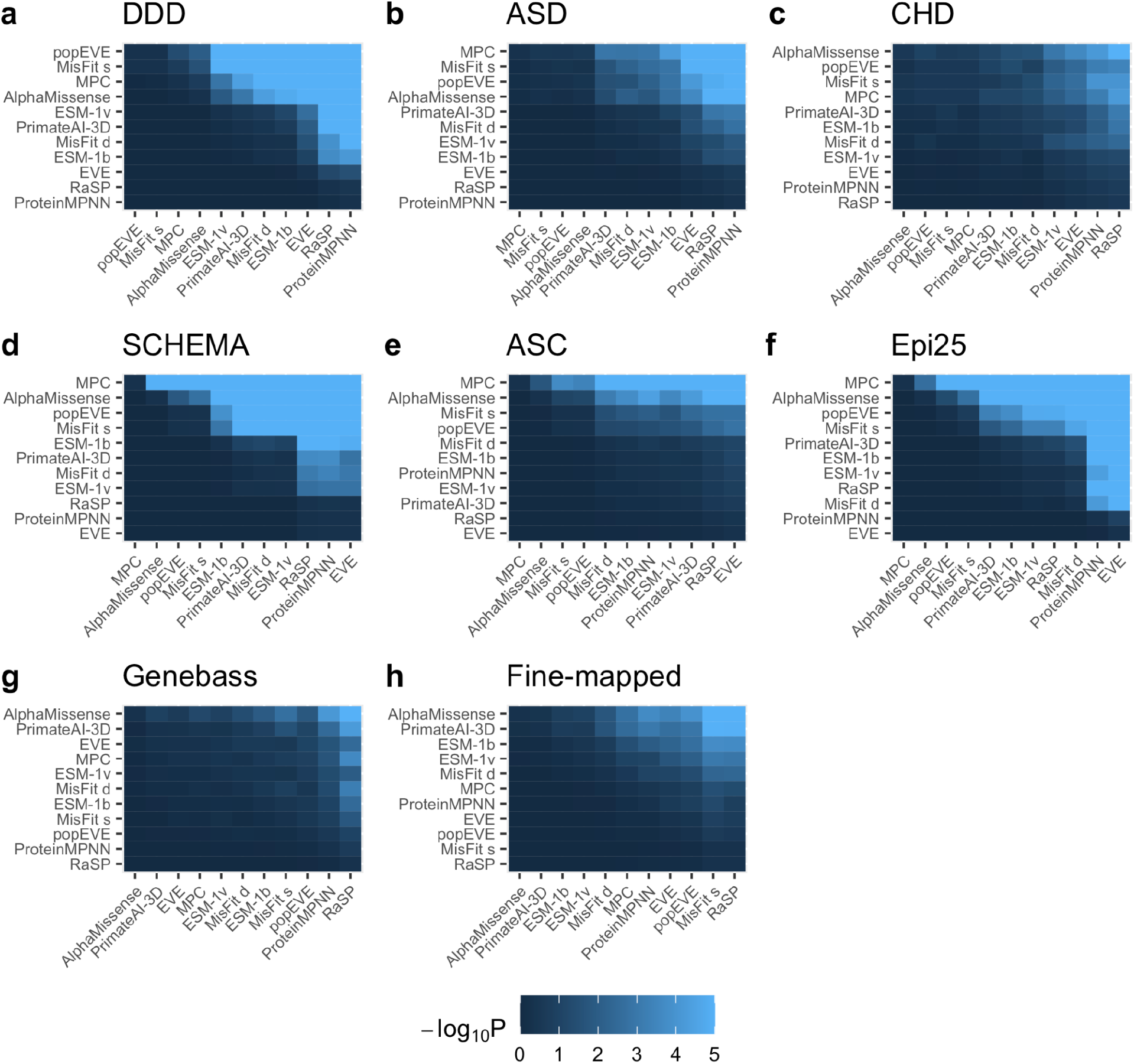
Significance of pairwise comparisons in Extended Data Figure 2 at a top 5% cutoff. For each evaluation data set in Extended Data Figure 2, for each pair of VSMs, the one-sided p-value assessing whether VSM 1 (row) outperforms VSM 2 (column), computed using the Fisher’s exact test.

**Extended Data Figure 4.**
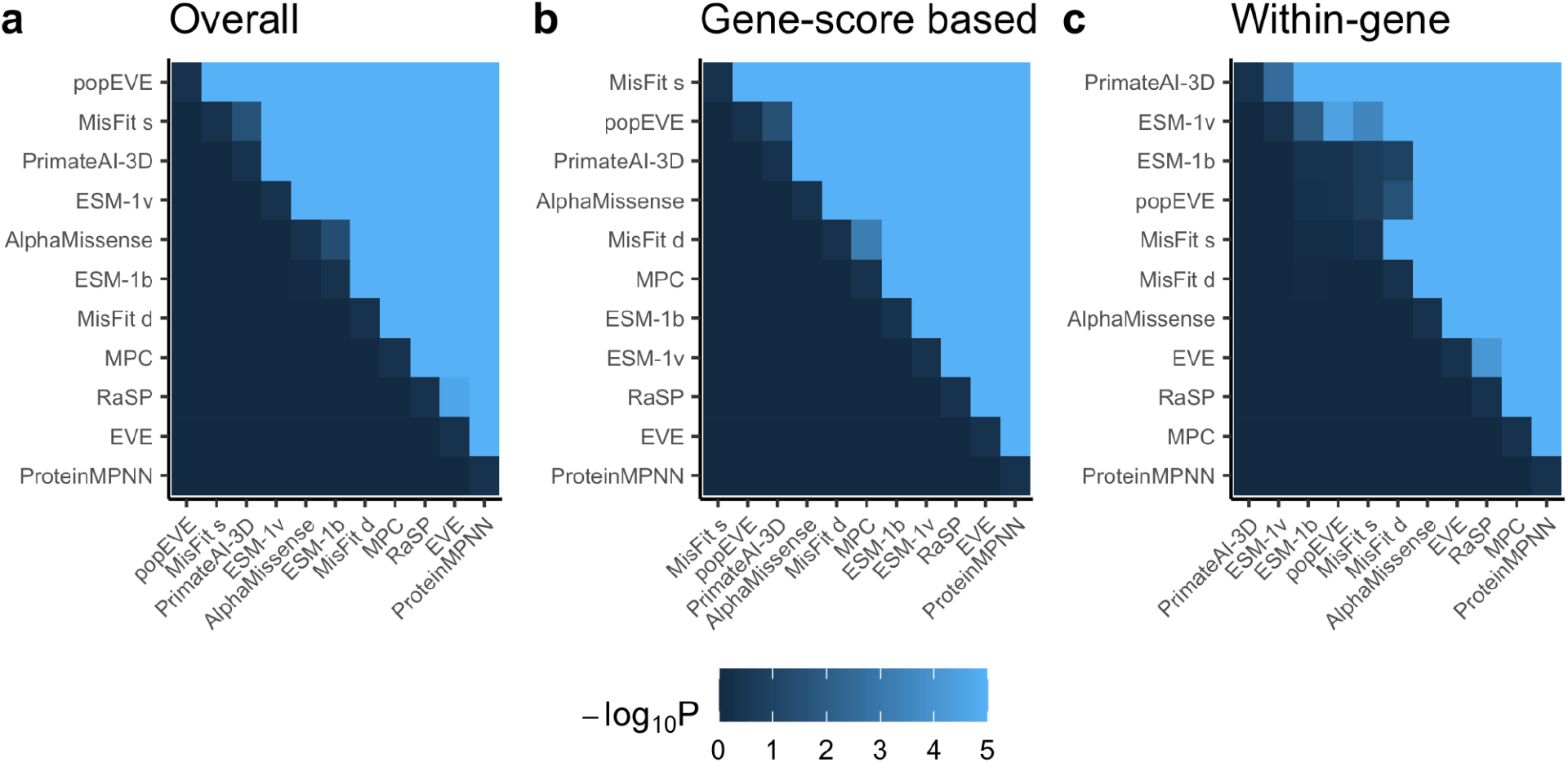
Significance of pairwise comparisons in Figure 2. For each evaluation data set in Figure 2, for each pair of VSMs, the one-sided p-value assessing whether VSM 1 (row) outperforms VSM 2 (column), computed using the Fisher’s exact test.

**Extended Data Figure 5.**
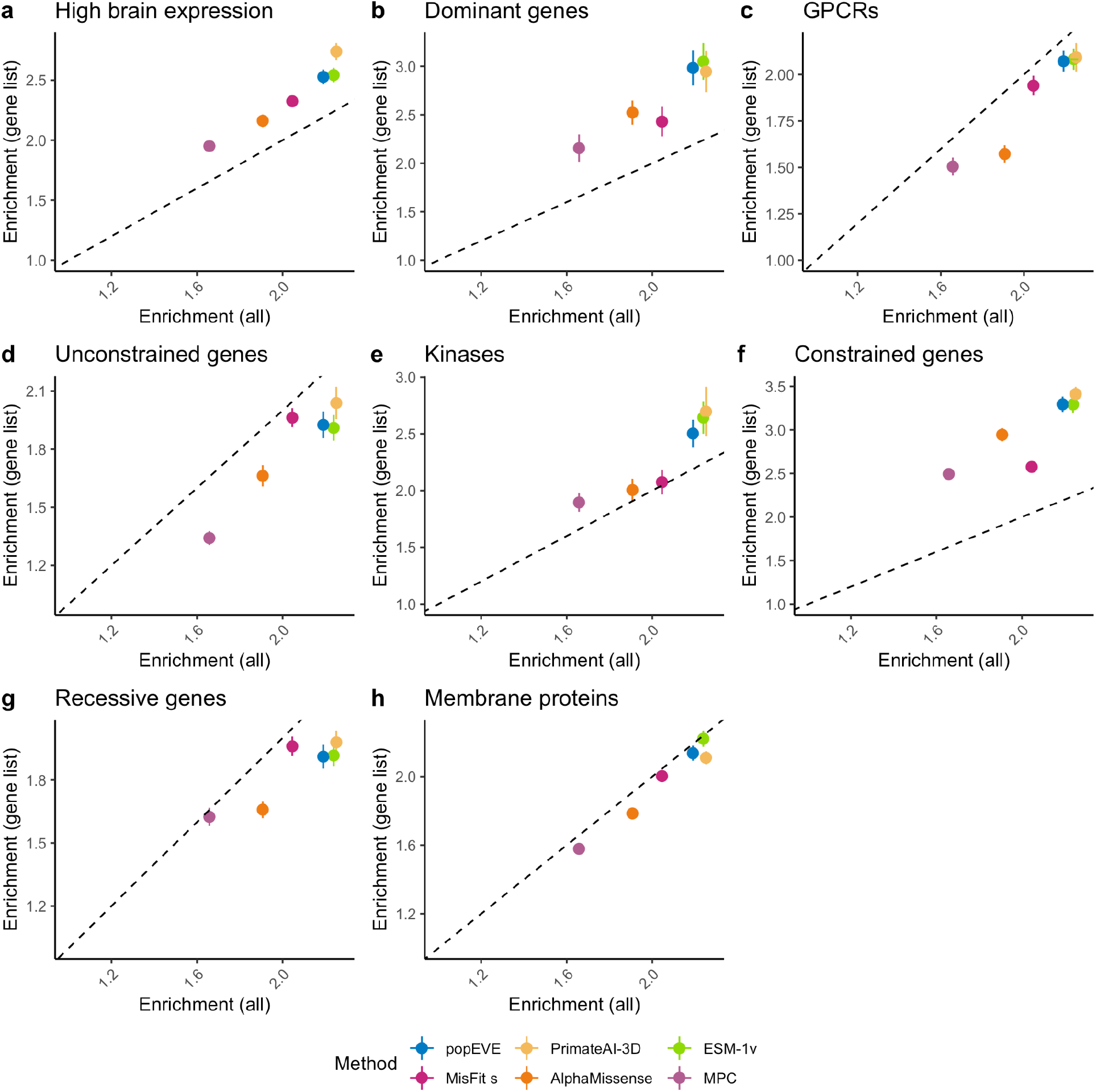
Evaluation of within-gene performance on subsets of genes, using gnomAD-new. Within-gene performance is computing by taking the average across genes of the enrichment of each VSM for identifying variants unobserved in gnomAD v4 vs variants observed in gnomAD-new, computed separately for each gene. Each panel compares the average within-gene enrichment computed within a subset of genes to the average within-gene enrichment computed on all genes. Gene subsets are as follows: (a) Top 10% of genes by brain expression in GTEx. (b) Dominant disease genes. (c) G-coupled protein receptors. (d) Top 10% of genes by LOEUF score. (e) Kinases. (f) Bottom 10% of genes by LOEUF score. (g) Recessive disease genes. (h) Cell surface proteins from the in silico human surfaceome^19^.

**Extended Data Figure 6.**
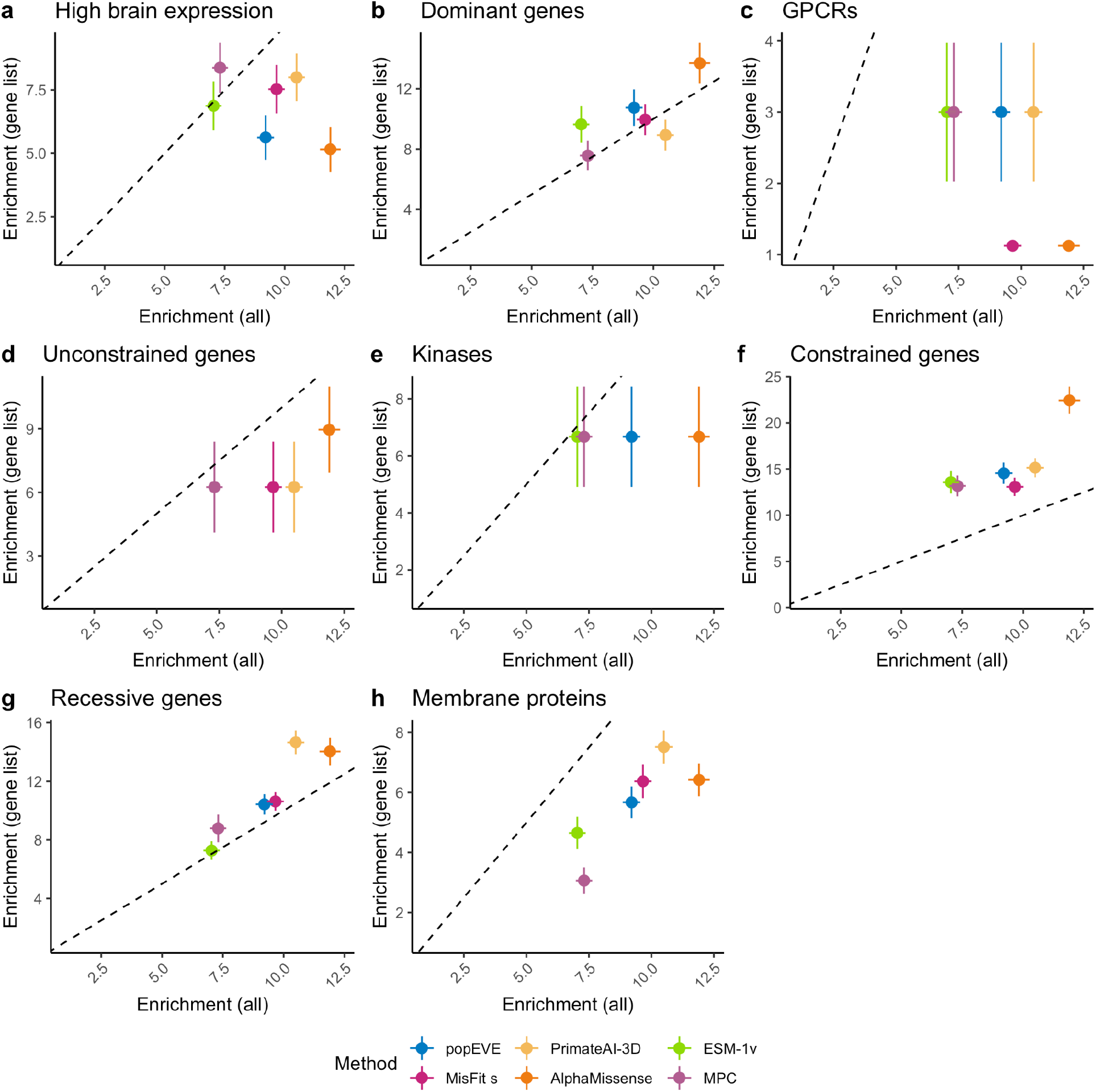
Evaluation of within-gene performance on subsets of genes, using ClinVar. Within-gene performance is computing by taking the average across genes of the enrichment of each VSM for identifying unobserved variants vs observed variants in ClinVar, computed separately for each gene. Each panel compares the average within-gene enrichment computed within a subset of genes to the average within-gene enrichment computed on all genes. Gene subsets are as follows: (a) Top 10% of genes by brain expression in GTEx. (b) Dominant disease genes. (c) G-coupled protein receptors. (d) Top 10% of genes by LOEUF score. (e) Kinases. (f) Bottom 10% of genes by LOEUF score. (g) Recessive disease genes. (h) Cell surface proteins from the in silico human surfaceome^19^.

## References

1. Adzhubei, I. A. et al. A method and server for predicting damaging missense mutations. Nat. Methods 7, 248–249 (2010).

2. Notin, P. et al. ProteinGym: Large-Scale Benchmarks for Protein Design and Fitness Prediction. bioRxiv (2023) doi:10.1101/2023.12.07.570727.

3. Livesey, B. J. & Marsh, J. A. Updated benchmarking of variant effect predictors using deep mutational scanning. Mol. Syst. Biol. 19, e11474 (2023).

4. Livesey, B. J. & Marsh, J. A. Variant effect predictor correlation with functional assays is reflective of clinical classification performance. bioRxiv 2024.05.12.593741 (2024) doi:10.1101/2024.05.12.593741.

5. Cheng, J. et al. Accurate proteome-wide missense variant effect prediction with AlphaMissense. Science 381, eadg7492 (2023).

6. Gao, H. et al. The landscape of tolerated genetic variation in humans and primates. Science 380, eabn8153 (2023).

7. Orenbuch, R. et al. Deep generative modeling of the human proteome reveals over a hundred novel genes involved in rare genetic disorders. medRxiv (2023) doi:10.1101/2023.11.27.23299062.

8. Chao, K. R. et al. The landscape of regional missense mutational intolerance quantified from 125,748 exomes. bioRxiv (2024) doi:10.1101/2024.04.11.588920.

9. Zhao, Y. et al. A probabilistic graphical model for estimating selection coefficient of missense variants from human population sequence data. medRxiv (2023) doi:10.1101/2023.12.11.23299809.

10. Rives, A. et al. Biological structure and function emerge from scaling unsupervised learning to 250 million protein sequences. Proc. Natl. Acad. Sci. U. S. A. 118, (2021).

11. Kaplanis, J. et al. Evidence for 28 genetic disorders discovered by combining healthcare and research data. Nature 586, 757–762 (2020).

12. Fu, J. M. et al. Rare coding variation provides insight into the genetic architecture and phenotypic context of autism. Nat. Genet. 54, 1320–1331 (2022).

13. Jin, S. C. et al. Contribution of rare inherited and de novo variants in 2,871 congenital heart disease probands. Nat. Genet. 49, 1593–1601 (2017).

14. Singh, T. et al. Rare coding variants in ten genes confer substantial risk for schizophrenia. Nature 604, 509–516 (2022).

15. Karczewski, K. J. et al. Systematic single-variant and gene-based association testing of thousands of phenotypes in 394,841 UK Biobank exomes. Cell Genomics 2, 100168 (2022).

16. Kanai, M. et al. Insights from complex trait fine-mapping across diverse populations. bioRxiv (2021) doi:10.1101/2021.09.03.21262975.

17. Satterstrom, F. K. et al. Large-Scale Exome Sequencing Study Implicates Both Developmental and Functional Changes in the Neurobiology of Autism. Cell 180, 568–584.e23 (2020).

18. Epi25 Collaborative, Chen, S., Neale, B. M. & Berkovic, S. F. Shared and distinct ultra-rare genetic risk for diverse epilepsies: A whole-exome sequencing study of 54,423 individuals across multiple genetic ancestries. medRxiv (2023) doi:10.1101/2023.02.22.23286310.

19. Bausch-Fluck, D. et al. The in silico human surfaceome. Proc. Natl. Acad. Sci. U. S. A. 115, E10988–E10997 (2018).

