## Supplementary Information for "Variant scoring performance across selection regimes depends on variant-to-gene and gene-to-disease components"

### GeneticsGym Supplementary Information

#### Evaluation data sources

We aggregated a series of datasets to benchmark variant scoring methods (VSMs), and we describe some considerations for each below. We combined the datasets using the Linker file (final section), based on available links: we used genetic variants where already mapped (locus and alleles), and Uniprot ID and amino acid position otherwise. For most data sets, we are not claiming that every mutation in the positive set is truly causal, but rather that the positive set is enriched over the negative set for disease-causing variants, and that the ability to distinguish these two sets is a worthwhile benchmark for a VSM.

We make processed evaluation tables available as hail tables (hail.is) and .tsv files in gs://genetics-gym/evaluation_tables/. All processed evaluation tables are in GRC38. For the largest files, the .tsv files do not include meta-data; for information such as gene names, please use the hail table or merge with a file that contains that information.

##### *de novo* variants

**Description:** Published datasets of *de novo* variation in trio-based studies of developmental delay (DD), autism spectrum disorder (ASD), and congenital heart disease (CHD) provide information about the high selection regime of variants.
**DD cases:** 26,400 *de novo* variants from 31,058 affected probands from (Kaplanis et al. 2020).
**ASD cases**: 11,077 *de novo* variants from 15,036 affected probands from (Fu et al. 2022).
**CHD cases:** 2,979 *de novo* variants from 2,871 affected probands from (Jin et al. 2017).
**Shared control set:** 5,563 *de novo* variants from 7,281 individuals, including 5,492 unaffected siblings from (Fu et al. 2022) + 1,789 controls from (Jin et al. 2017).
**Raw files:**

- DD (probands): gs://genetics-gym/raw_data/kaplanis_variants_annotated_2024-05-15.txt
- ASD (probands and unaffected siblings): gs:/genetics-gym/raw_data/Nat_Gen_published_autosomal_and_updated_XY_de_novo_calls_2024-05-13_no_cohort.txt
- CHD (probands): gs://genetics-gym/raw_data/NIHMS906719-chd-dnm-cases.csv
- Controls (from CHD study): gs://genetics-gym/raw_data/NIHMS906719-chd-dnm-controls.csv

**Processed files:**

- DD: gs://genetics-gym/evaluation_tables/dd-with-combined-controls{.ht, .tsv.bgz)
- ASD: gs://genetics-gym/evaluation_tables/asd-with-combined-controls{.ht, .tsv.bgz)
- CHD: gs://genetics-gym/evaluation_tables/chd-with-combined-controls{.ht, .tsv.bgz)

#### Rare variant associations

**Description:** Previously generated rare variant association data from case-control studies of schizophrenia (SCHEMA), autism (ASC), and epilepsy (Epi25) identify rare variants as more likely to influence these disorders.
**Positive set:** Variants found in at least one case, but not in controls (820,282 SCHEMA, 246,016 ASC, 1,235,640 Epi25).
**Negative set:** Variants found in at least one control, but not in cases (3,793,987 SCHEMA, 435,914 ASC, 2,476,453 Epi25).
**Raw files:**

- SCHEMA: gs://genetics-gym/raw_data/SCHEMA_variant_results.vcf.bgz
- ASC: gs://genetics-gym/raw_data/ASC_variant_results.vcf.bgz
- Epi25: gs://exome-results-browsers-public/downloads/2022-12-01/Epi25/Epi25_variant_results.vcf.bgz

**Processed files:**

- SCHEMA: gs://genetics-gym/evaluation_tables/schema_evaluation_table{.ht, .tsv.bgz)
- ASC: gs://genetics-gym/evaluation_tables/asc_evaluation_table{.ht, .tsv.bgz)
- Epi25: gs://genetics-gym/evaluation_tables/epi25_evaluation_table{.ht, .tsv.bgz)

#### Biobank fine-mapping

**Description:** We previously performed statistical fine-mapping in UK Biobank, Biobank Japan, and FinnGen to identify likely causal variants for complex traits and common diseases (Kanai et al. 2021). Because fine-mapping is only powered for common variants, we match our positive and negative sets on minor allele frequency.
**Positive set:** 1,688 variants with PIP > 0.1 for at least one phenotype in at least one biobank.
**Negative set:** 50,671 variants with PIP < 0.01, matched to the positive set on MAF.
*Note: Because these files rely on data from FinnGen that is not yet public, we are not able to release them at this time. These are slated to be public by the end of 2024, at which point we will provide this file.*

#### Biobank rare variants

**Description:** We have previously performed a systematic rare variant association analysis of 4,529 phenotypes across the UK Biobank (Karczewski et al. 2022). There, we showed that after adjusting for power as governed by allele frequency, we observe a higher proportion of pLoF variants associated with at least one trait, compared to missense variants, compared to synonymous variants. We use a similar principle to score genetic variants based on impact.
**Positive set:** 12,510 variants associated with at least one trait with high confidence.
**Negative set:** 12,497 variants not associated with any trait; matched to the positive set on MAF.
**Raw files:** gs://genetics-gym/raw_data/mutation/genebass.ht
**Processed file:** gs://genetics-gym/evaluation_tables/genebass_evaluation_table{.ht, .tsv.bgz)

##### Natural variation

**Description:** The VSMs benchmarked here were trained before gnomAD v4 data was available. However, several methods made use of gnomAD v2 and/or UK Biobank data, which is included in gnomAD v4. To conduct an independent benchmarking, we excluded any variants observed in gnomAD v2 and/or UK Biobank from the entire analysis.
**Positive set:** 83,260,797 variants not observed in gnomAD v4
**Negative set:** 13,450,942 variants not observed in gnomAD v2 or in UK Biobank exome data, but observed in gnomAD v4
**Raw files:**

- gs://gcp-public-data--gnomad/release/4.1/ht/exomes/gnomad.exomes.v4.1.sites.ht
- gs://gcp-public-data--gnomad/release/2.1.1/liftover_grch38/ht/exomes/gnomad.exomes.r2.1.1.sites.liftover_grch38.ht
- gs://ukbb-exome-public/500k/results/vep.ht

**Processed file:** gs://genetics-gym/evaluation_tables/gnomad-independent-set{.ht, .tsv.bgz}

##### ClinVar

**Description:** As variants deposited into ClinVar have biases that are hard to characterize and some VSMs are trained on ClinVar variation (explicitly, or implicitly by way of standing variation that is often characterized as benign), we do not recommend these as a frontline assessment. However, we use these data to compare performance across different gene classes.
**Positive set:** 235,772 Pathogenic or Likely Pathogenic variants
**Negative set:** 1,110,947 Benign or Likely Benign variants
**Raw files:** https://ftp.ncbi.nlm.nih.gov/pub/clinvar/vcf_GRCh38/clinvar.vcf.gz (downloaded on 4/25/24)

##### Linker file

**Description:** To aggregate the different methods and compare them on the same variants, we created a linker table based on VEP GRCh38 for the autosomes and the X chromosome. We computed the most severe consequence using https://broadinstitute.github.io/gnomad_methods/api_reference/utils/vep.html#gnomad.utils.vep.process_consequences. Each variant is keyed by locus and allele and annotated with ensembl transcript ID, ensembl gene ID, uniprot ID and HGNC gene ID.
**Raw files:**
gs://gcp-public-data--gnomad/resources/context/grch38_context_vep_annotated.v105.ht/
**Processed file:** gs://genetics-gym/linkers-and-annotations/linker_no_mane_filter.ht

#### Gene annotations

For gene list specific assessments in Figure 2 and Extended Data Figures 5-6, we restricted analysis to genes in particular gene sets including:

- Brain genes: https://storage.googleapis.com/adult-gtex/bulk-gex/v8/rna-seq/GTEx_Analysis_2017-06-05_v8_RNASeQCv1.1.9_gene_median_tpm.gct.gz - top 10% for brain.
- Recessive, dominant, kinases, and GPCRs downloaded from https://github.com/macarthur-lab/gene_lists
- Constrained genes: top and bottom deciles by loeuf from gs://gcp-public-data--gnomad/release/4.0/constraint/gnomad.v4.0.constraint_metrics.ht
- Membrane proteins downloaded from https://wlab.ethz.ch/surfaceome/surfaceome_ids.txt

### Evaluations and significance

**Enrichments and rate ratios.**
For a given VSM *v*, threshold *T*, and evaluation data set with label *y*, let

- TP = |{*i* : *v*(*i*)>*T* and *y*(*i*)=1}|
- FP = |{*i* : *v*(*i*)>*T* and *y*(*i*)=0}|
- FN = |{*i* : *v*(*i*)<*T* and *y*(*i*)=1}|
- TN = |{*i* : *v*(*i*)<*T* and *y*(*i*)=0}|

We define enrichment as (TP / (TP + FN)) / (FP / (FP + TN)). For a data set of de novo variants in N1 probands and N2 controls, we define rate ratio as (TP / N1) / (FP / N2).

Different VSMs have scores available for different sets of possible mutations, which may affect performance metrics in unpredictable ways. For analyses with gnomAD or ClinVar data, we restricted to the set of potential mutations that was scored by all VSMs. For other data sets, however, this resulted in a prohibitive decrease in the number of potential mutations available for evaluation. For the DD, ASD, CHD, Fine-mapped, Genebass, SCHEMA, ASC, and Epi25 data sets, we sought to create an analytic approach that maximized the number of variants analyzed while providing a fair comparison (**Box 1**).

Suppose we are analyzing VSMs $V_1, \dots, V_K$ with scores on sets $S_1, \dots S_K$ of potential mutations, respectively. Let $e(V, S)$ denote the enrichment of VSM $V$ on set $S$ of variants. We can compute $e(V_i, S_i)$, for each $i$, but these may not be comparable to each other. Instead, we assume that ratios of these enrichments are not too sensitive to the set of variants on which they are computed. To leverage this assumption, We let $V^*$ denote the VSM defined on the largest set of variants, $S^*$. We then estimate the enrichment of each VSM on the set $S^*$ by

$$\widehat{enr}(V_i, S^*) := enr(V^*, S^*) \frac{enr(V_i, S_i \cap S^*)}{enr(V^*, S_i \cap S^*)}.$$

We use the bootstrap to compute standard errors. The computation for rate ratios is analogous. For each pairwise comparison, the set of variants that had a positive or negative label in the evaluation set and that was scored by both methods was the baseline for computing the percentile thresholds.

**Box 1**. Description of the method for estimating enrichments and rate ratios for sets of VSMs that do not have scores available on a large enough set of overlapping mutations.

**Significance.** We computed significance on the pairwise comparisons among all methods. For simplicity, we chose a single percentile threshold, 0.95, at which to compute significance. We used a different approach for within-gene vs genome-wide evaluations

- Genome-wide comparisons: To compare two VSMs, we restricted to the set of sites at which the evaluation labels and both VSMs were defined. We then used Fisher's Exact Test to determine whether sites with percentile($VSM_1$) > 0.95 or sites with percentile($VSM_2$) > 0.95 were significantly more likely to have a positive evaluation label.
- Within-gene comparisons: For each gene, we recorded whether $VSM_1$ outperformed $VSM_2$ within that gene. To do this, we computed within-gene percentiles for $VSM_1$ and $VSM_2$, and we assessed whether sites with percentile($VSM_1$) > 0.95 or sites with percentile($VSM_2$) > 0.95 were more likely to have a positive evaluation label. We then performed a binomial test of whether $VSM_1$ outperformed $VSM_2$ significantly more than 50% of the time.

### Variant Scoring Methods

We downloaded publicly available scores for ESM1b, ESM1v, MisFit, MPC, PopEVE, EVE, and RaSP. We obtained PrimateAI-3D scores with permission from Illumina. We computed ProteinMPNN Log Likelihood Ratios using AlphaFold predicted structures for the human proteome.

### References

Fu, Jack M., F. Kyle Satterstrom, Minshi Peng, Harrison Brand, Ryan L. Collins, Shan Dong, Brie Wamsley, et al. 2022. “Rare Coding Variation Provides Insight into the Genetic Architecture and Phenotypic Context of Autism.” *Nature Genetics* 54 (9): 1320–31.

Jin, S. C., J. Homsy, S. Zaidi, Q. Lu, S. Morton, S. R. DePalma, X. Zeng, et al. 2017. “Contribution of Rare Inherited and de Novo Variants in 2,871 Congenital Heart Disease Probands.” *Nature Genetics* 49 (11). https://doi.org/10.1038/ng.3970.

Kanai, Masahiro, Jacob C. Ulirsch, Juha Karjalainen, Mitja Kurki, Konrad J. Karczewski, Eric Fauman, Qingbo S. Wang, et al. 2021. “Insights from Complex Trait Fine-Mapping across Diverse Populations.” *bioRxiv*. https://doi.org/10.1101/2021.09.03.21262975.

Kaplanis, Joanna, Kaitlin E. Samocha, Laurens Wiel, Zhancheng Zhang, Kevin J. Arvai, Ruth Y. Eberhardt, Giuseppe Gallone, et al. 2020. “Evidence for 28 Genetic Disorders Discovered by Combining Healthcare and Research Data.” *Nature* 586 (7831): 757–62.

Karczewski, Konrad J., Matthew Solomonson, Katherine R. Chao, Julia K. Goodrich, Grace Tiao, Wenhan Lu, Bridget M. Riley-Gillis, et al. 2022. “Systematic Single-Variant and Gene-Based Association Testing of Thousands of Phenotypes in 394,841 UK Biobank Exomes.” *Cell Genomics* 2 (9): 100168.
